# Phasor diagrams clock oscillatory hemodynamic switching between overt speech production and micro resting states

**DOI:** 10.1101/2025.07.29.667380

**Authors:** Ruey-Song Huang, Mengxiao Wang, Teng Ieng Leong, Ut Meng Lei, Martin I. Sereno, Defeng Li, Victoria Lai Cheng Lei

## Abstract

This study investigates the intricate interplay between the task-positive network and the default mode network (DMN) during transitions between overt language tasks and brief resting periods. While previous research suggests that these networks are not invariably anticorrelated, the precise timing of transitions has remained elusive. We employed rapid phase-encoded fMRI to decode brain dynamics with ultimate precision, capturing these transitions in real time. By utilizing phasor diagrams to represent the oscillatory activities, we examined the amplitudes and phases of hemodynamic fluctuations within the language network and DMN. Our findings align with existing empirical and theoretical perspectives on DMN functions and cognitive task performance, affirming the validity of our approach. We identified heterogeneous micro resting states interwoven with periods of overt speech production. Notably, various core regions of the DMN exhibited task-dependent amplitude and phase modulations, with activation strength and delay rising in line with increasing task complexity, ranging from comprehension to immediate and delayed speech production. This study sheds light on the dynamic engagement of the DMN during overt speech production, providing precise timing data of transitions between the DMN and language network. It demonstrates that rapid phase-encoded fMRI and phasor diagrams are powerful tools for measuring the switching between active tasks and micro resting states with subsecond accuracy, while also elucidating task load-dependent changes in the DMN. By accurately measuring the timing of these transitions, we gain insights into cognitive flexibility, attention, and the efficiency of information processing.

## Introduction

Naturalistic tasks, such as overt speech production, involve frequent transitions between action and rest. Functional magnetic resonance imaging (fMRI) studies have revealed an anticorrelation between the task-positive network (TPN) and task-negative default mode network (DMN) (1). These alternating activations between the TPN and DMN occur both during continuous resting periods (1-3) and in experiments comprising task and rest periods (3-6). The overall DMN activations during intermittent resting periods in blocked-design experiments resemble those observed in continuous resting-state (3, 5, 6). In event-related design experiments, however, the timing of DMN activities varies depending on the task phases and load (7, 8). While lagged correlation analysis has revealed the latency structure of the TPN and DMN in continuous resting-state fMRI data (9), how DMN activities are dynamically modulated by the alternation between overt speech production and rest remains unclear. Here, we unravel the event-related dynamics of the TPN and DMN during language perception and production tasks using rapid phase-encoded fMRI designs (10, 11). We applied phasor diagrams, graphical representations of oscillatory quantities traditionally used in physics and engineering, to characterize the amplitudes and phases of periodic hemodynamic fluctuations in the TPN and DMN, as well as the dynamic switching between them. We aim to identify task-dependent modulations in the DMN and its role in overt language production.

## Results

This section presents the results from an fMRI experiment involving language perception and production in Chinese (L1) and English (L2) (*Supporting Information*). Fig. 1A shows significant periodic activations (F(2,230) > 12.11, P < 10-5, uncorrected) on the cortical surface of Subject 1 during an English reading-reciting task. Fig. 1B depicts periodic activations with progressive delays in five surface-based regions of interest (sROIs). These time courses are color-coded by activation phases, which are visualized in a phasor diagram (Fig. 1C and D).

**Figure 1.**
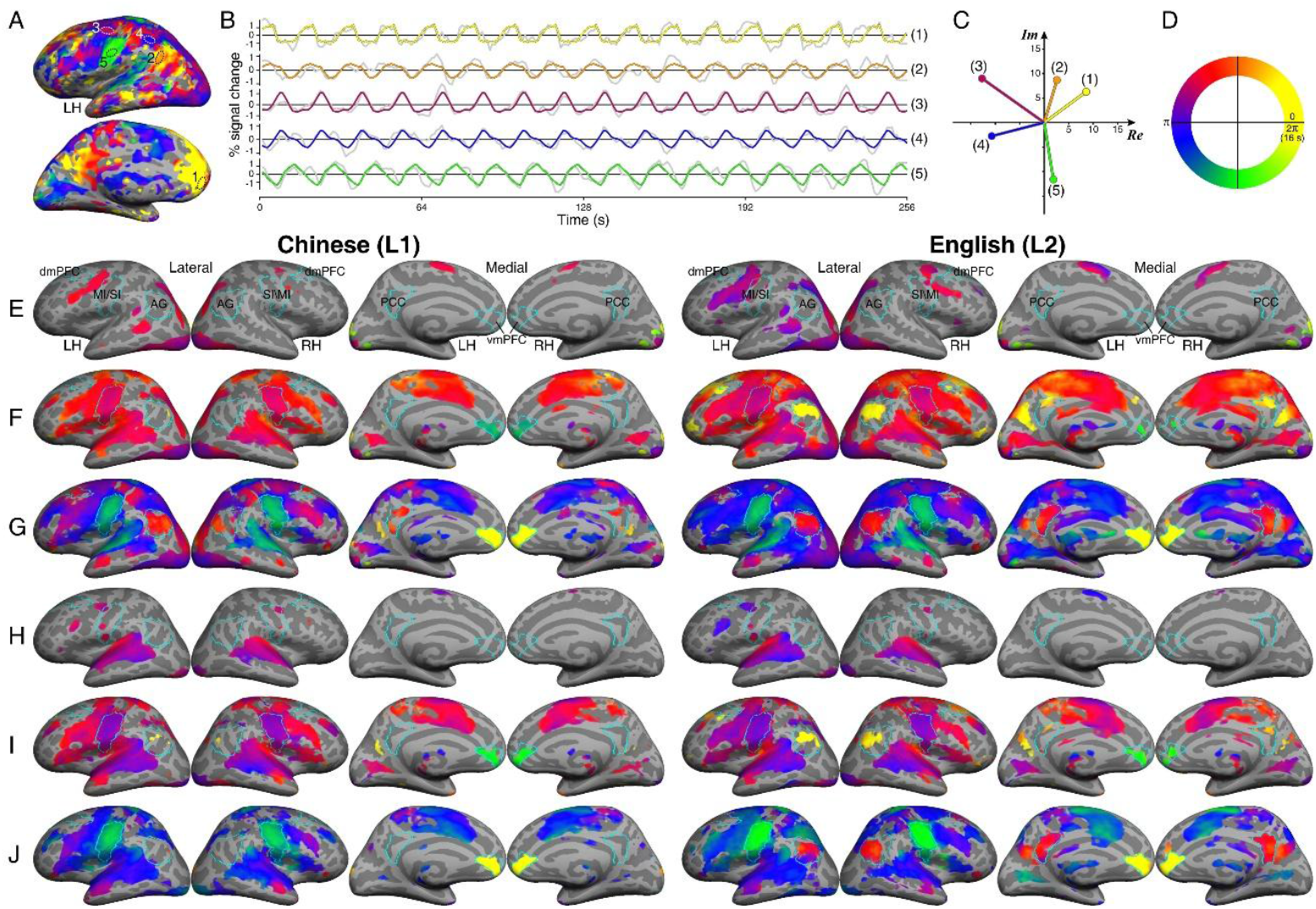
(A) A phase-encoded activation map of Subject 1. (B) Average time courses within five sROIs outlined in (A). Gray curves: time courses averaged across voxels in each sROI; Colored curves: 16 repetitions of the average time course within a 16-s period. (C) A phasor diagram showing the average complex F-values in each sROI. *Re*: real, *Im*: imaginary. (D) Color-coded activation phases in [0^°^, 360^°^] or [0 s, 16 s]. (E-J) Group-average activation maps (N=31) for six tasks: silent-reading (E), reading-aloud (F), reading-reciting (G), listening (H), shadowing (I), and listening-reciting (J). Statistical significance: *F*(2,230) > 4.7, *P* < 0.01 (subject level); *F*(2, 60) > 4.97, *P* < 0.05, cluster-corrected (group level). LH: left hemisphere; RH: left hemisphere; Cyan contours: sROIs in the DMN and MI/SI.

Fig. 1E-J shows group-average activation maps for six tasks performed in either Chinese (L1) or English (L2). The silent-reading task activated areas in the posterior parietal, ventral occipitotemporal, and temporal cortices (Fig. 1E), while the listening task engaged the superior temporal cortex (Fig. 1H). Both language comprehension tasks also activated areas in the premotor and supplementary motor cortices. The other four tasks, which included overt speech production, induced further activations in the primary motor and somatosensory (MI/SI orofacial regions), cingulate motor, prefrontal, and insular cortices (Fig. 1F, G, I, and J). Overall, L1 and L2 maps show the same activation patterns across tasks, exhibiting larger extent in L2 tasks involving speech production. The DMN does not show significant activations in comprehension-only tasks, even though they included an 11-s rest per trial (Fig. 1E, H). Most of the DMN core regions, including dorsomedial prefrontal cortex (dmPFC), ventromedial prefrontal cortex (vmPFC), angular gyrus (AG), and posterior cingulate cortex (PCC), were activated in all tasks involving overt speech production (Fig. 1F, G, I, and J).

For each task in each language, two phasor diagrams (left and right hemispheres; Fig. 2A) were created to compare the activation amplitudes and phases in task-positive (“Task”) regions and four DMN core regions (Fig. 1E). The silent-reading and listening tasks induced significant activations in Task regions, with comparable phases across tasks (blue phasors). Among DMN core regions, only vmPFC shows consistent activation phases in both tasks (red phasors). The activation amplitudes in vmPFC increased significantly in the reading-aloud and shadowing tasks, both involving immediate overt speech output. In these two tasks, the average activation phases in vmPFC (red phasors) are delayed from those in Task regions (blue phasors) by 124.63^°^ ± 12.64^°^ (or 5.54 ± 0.56 s) in the counterclockwise direction (Dataset S1).The activation phases in Task regions during the reading-reciting and listening-reciting tasks, both involving working memory and overt speech output, exhibit a 59.36^°^ ± 3.47^°^ (or 2.64 ± 0.15 s) delay from those in Task regions during the reading-aloud and shadowing tasks. In the reading-reciting and listening-reciting tasks, the activation phases in vmPFC differ from those in Task regions by 178.01^°^ ± 14.88^°^ (or 7.91 ± 0.66 s), in approximately antiphase to each other. Furthermore, the activation phases in vmPFC show a 122.31^°^ ± 14.73^°^ (or 5.44 ± 0.65 s) delay from those in the MI/SI orofacial region (black needles in Fig. 2A) in all tasks involving overt speech production. Across all L2 tasks, the activation phases in Task regions, vmPFC, and MI/SI in the left hemisphere exhibit a slight delay of 13.30^°^ ± 9.32^°^ (or 0.59 ± 0.41 s) from those in L1 tasks.

**Figure 2.**
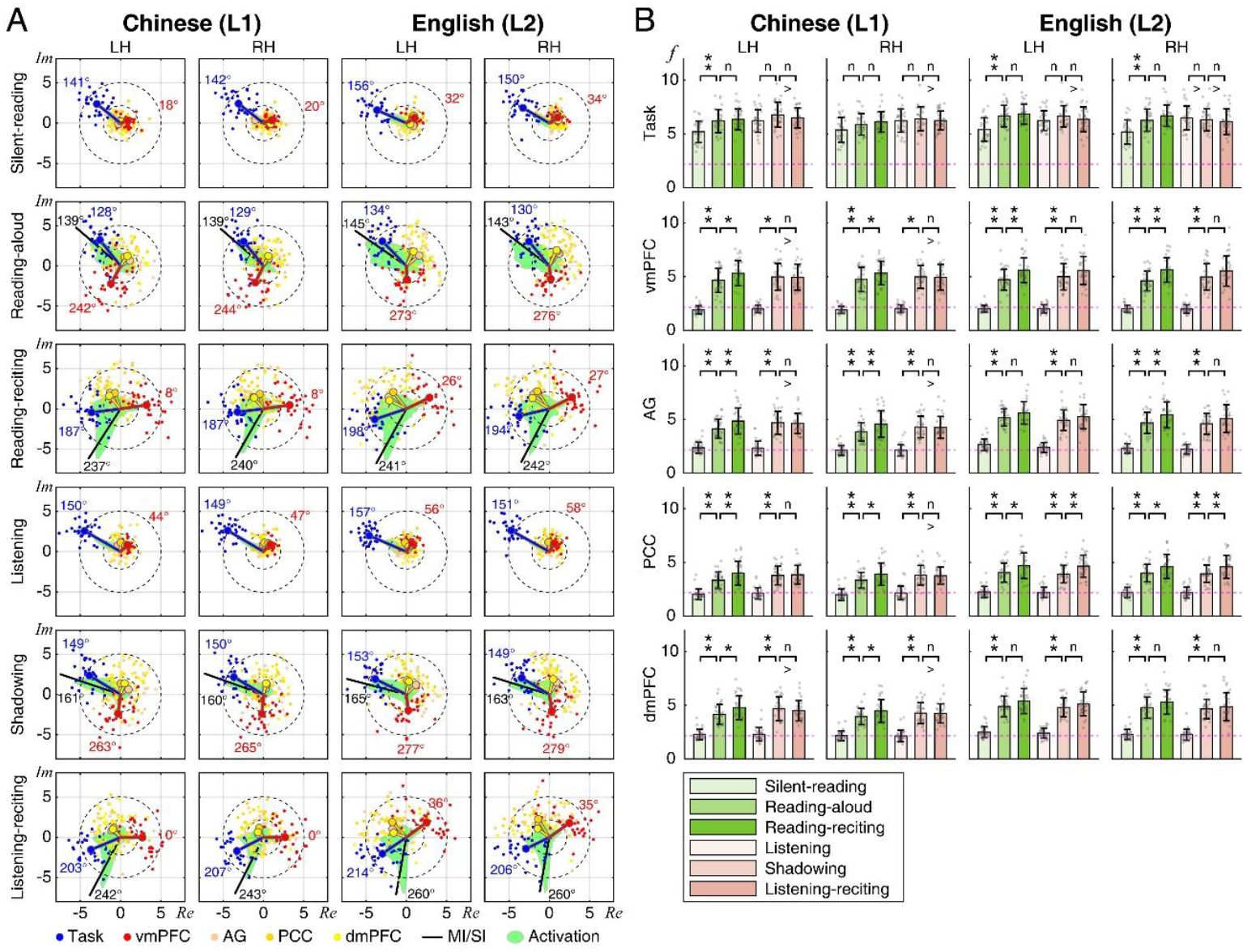
(A) Phasor diagrams. Lollipop or needle: an average complex *F*-value of vertices within a Task region or an sROI in the group-average maps (Fig. 1E-J; *Supporting Information*). Colored dot: an average complex *F*-value within a Task region or an sROI in the single-subject maps. *Re*: real, *Im*: imaginary. Inner dashed circle: *F*(2,230) = 4.7, *P* = 0.01, uncorrected; Outer dashed circle: *F*(2,230) = 25.5, *P* = 10^−10^, uncorrected. (B) Cross-subject distribution of average activation amplitude (*f*) in each sROI (*N* = 31; error bar: standard deviation; *Supporting Information*). Magenta dashed line: *F*(2,230) = 4.7, *P* = 0.01, uncorrected. Bracket: a paired *t*-test between tasks, ** *P* < 0.01, * *P* < 0.05, corrected; n: not significant (*P* > 0.05); > sign: the left bar is higher than the right bar.

Fig. 2B shows the distributions of activation amplitudes in single-subject maps for each task (*Supporting Information*). The inter-subject average amplitudes in all DMN sROIs in the left hemisphere exhibit a staircase pattern in line with increasing task complexity, ranging from comprehension-only to immediate production to delayed production in both languages.

## Discussion

Inspired by the three-phase alternating current system for power distribution, we used phasor diagrams to characterize periodic activations in the TPN and DMN as well as alternations between them with subsecond precision. We found that the DMN was dynamically modulated by tasks involving overt speech production, but not by comprehension-only tasks. Among the DMN core regions, vmPFC exhibited the strongest task-dependent modulations, with activation amplitudes increasing from comprehension to immediate and delayed production tasks. The activation phases in vmPFC differed from those in Task regions by ∼120^°^ and ∼180^°^ in immediate and delayed production tasks, respectively. However, vmPFC consistently shows a ∼120^°^ delay from the MI/SI orofacial region in all tasks involving production. These findings suggest that the DMN is not always in anticorrelation (i.e., 180^°^ out of phase) with the TPN, and that vmPFC is highly coupled with the overt production phase. Furthermore, vmPFC showed larger and later activations in second language tasks, reflecting a dependency on task nature (7-8, 12-13).

Other DMN core regions, including AG, PCC, and dmPFC, exhibited different activation patterns. Their activation phases fell within the task-positive periods, suggesting they were engaged in sentence-level semantic encoding for immediate production or silent rehearsal (inner speech) for delayed production. These regions have been suggested to be involved in self-referential processing and self-monitoring during overt speech production (4, 12-14). Together, phase-encoded fMRI revealed multiple heterogeneous micro resting states interleaving periods of overt speech production. Different DMN core regions showed task-dependent modulations at multiple phases, suggesting dynamic engagements of the DMN during language production (7, 13-15).

In conclusion, the application of phasor diagrams for visualizing multi-phase brain activities represents a promising advancement in cognitive neuroscience. By decoding amplitude and phase modulations within the TPN and DMN, we can now better understand and quantify the intricate dynamics of cognitive processes. This method not only aids in assessing cognitive load and flexibility but also offers a novel perspective on the brain’s ability to transition seamlessly between active cognitive tasks and micro resting states. Such insights are critical for enhancing our understanding of neural efficiency and cognitive performance. This approach, therefore, holds significant potential for future research aimed at unraveling the complexities of various cognitive functions besides language production.

## Materials and Methods

### Methods appear in *Supporting Information*

#### Ethical Note

All subjects gave written informed consent in accordance with the protocols approved by the Research Ethics Committee of the University of Macau. All participants in this study were verbally informed of the nature and possible consequences of the study.

#### Data and Software Availability

Supplementary Dataset S1 is available on the Open Science Framework (https://osf.io/hea2p/?view_only=34d7f3c5a37341e9813e04796b0a25d3). Custom codes for analyzing phase-encoded fMRI data are included in *csurf* available at https://pages.ucsd.edu/~msereno/csurf/

## Acknowledgments

This research was supported by Macau Science and Technology Development Fund (FDCT 0001/2019/ASE), University of Macau Development Foundation (EXT-UMDF-014-2021), University of Macau (SRG2019-00189-ICI, MYRG2022-00265-ICI, MYRG2022-00200-FAH, CRG2020-00001-ICI, CRG2021-00001-ICI, CPG2023-00016-FAH, MYRG-CRG2024-00047-ICI), and National Institutes of Health (R01 MH081990 to M.I.S and R.S.H). We thank Yi Tang for participant recruitment, Cheok Teng Leong and Chi Un Choi for helping with fMRI experiments, and Yafang Li for image preprocessing.

## Supporting Information

### Methods

#### Participants

This study analyzed structural and functional images of 31 bilinguals (22 females, 9 males; mean age = 23.3 ± 4 years) from the Brain and Language Project at the University of Macau. The group-average maps of task-positive activations in 21 out of the 31 subjects in this study have been published elsewhere (Lei et al., 2024). All subjects were native Chinese (Mandarin) speakers who acquired English as a second language between the ages of 4 and 13 (mean age of acquisition = 7.1 ± 2.1 years). English proficiency was assessed based on their most recent standardized test results (IELTS score of 6.5 or above, or an equivalent qualification). All subjects gave written informed consent in accordance with the protocols approved by the Research Ethics Committee of the University of Macau. All of them had normal or corrected-to-normal vision and reported no history of neurological disorders.

#### Experimental Design and Setup

Each subject participated in one Chinese (L1) and one English (L2) fMRI session, each comprising two 256-s functional scans for each of the following sentence-level language processing tasks (Fig. 1): (1) silent reading for 5 s (Fig. 1E); (2) reading aloud for 5 s (Fig. 1F); (3) silent reading for 5 s, followed by reciting for 5 s (Fig. 1G); (4) listening for 5 s (Fig. 1H); (5) shadowing for 5 s (Fig. 1I); or (6) listening for 5 s, followed by reciting for 5 s (Fig. 1J). Each scan consisted of sixteen trials of task-rest alternation in a 16-s period, during which the resting duration was 11 s for tasks 1, 2, 4, and 5, and 6 s for tasks 3 and 6. A sentence with the subject-verb-object (SVO) structure was randomly selected and presented in each trial (see Lei, 2024 for details on language stimuli). In a 15-min practice session, subjects were trained to keep their heads still during speech production in an MRI simulator. In an fMRI session, subjects wore a thermoplastic mask and earplugs, and then put on MR-compatible noise-canceling headphones (OptoActive II, OptoAcoustics Ltd.), through which auditory stimuli and speech output were heard in real time (see Lei et al., 2024, for details on experimental setup).

#### Image Acquisition

Functional and structural brain images were acquired using a 32-channel head coil in a Siemens MAGNETOM Prisma 3T MRI scanner at the Centre for Cognitive and Brain Sciences, University of Macau. Each functional scan (64×64×55 voxels, 256 time points) was acquired using a blipped-CAIPIRINHA simultaneous multi-slice (SMS), single-shot echo-planar imaging (EPI) sequence with the following parameters: acceleration factor = 5; interleaved ascending slices; TR = 1000 ms; TE = 30 ms; flip angle = 60º; 55 axial slices; field of view = 192×192 mm; matrix size = 64×64; voxel size = 3×3×3 mm; 256 TR per image; dummy = 6 TR. Two sets of T1-weighted structural images were scanned using an MPRAGE sequence with the following parameters: TR = 2300 ms; TE = 2.26 ms; TI = 900 ms; flip angle = 8º; 256 axial slices; field of view = 256×256 mm; matrix size = 256×256; voxel size = 1×1×1 mm. The slice center and orientation of the structural images were the same as those of the functional images.

#### Image Preprocessing

Structural and functional images were analyzed by Analysis of Functional NeuroImages (*AFNI*; https://afni.nimh.nih.gov/), *FreeSurfer* (https://surfer.nmr.mgh.harvard.edu/), and *csurf* (https://pages.ucsd.edu/~msereno/csurf/) packages. Cortical surfaces were reconstructed from the structural images of each subject using *FreeSurfer* (see Lei et al., 2024, for details on data preprocessing).

#### Fourier-based Analysis

For each 4D functional dataset (64×64×55 voxels, 256 TR per voxel) of each task, the time series *x*_*m*_(*t*) of voxel *m* was analyzed using a 256-point discrete Fourier transform (Chen et al., 2019; Lei et al., 2024):

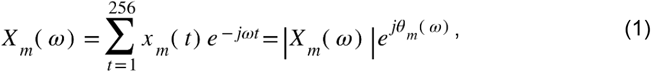

where *X*_*m*_(*ω*) is the complex number at frequency *ω* (0-127 cycles per scan), and |*X*_*m*_(*ω*)| and *θ*_*m*_(*ω*) are its amplitude and phase. *X*_*m*_(*ω*_*s*_) and *X*_*m*_(*ω*_*n*_) represent the periodic signal at task frequency *ω*_*s*_ (s = 16 cycles per scan) and noise at non-task frequencies *ω*_*n*_, respectively (Chen et al., 2019; Lei et al., 2024). The statistical significance of periodic activation in voxel *m* is evaluated as a signal-to-noise ratio (SNR):

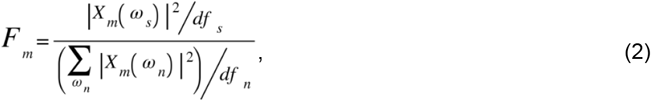

where *df*_*s*_ and *df*_*n*_ are the degrees of freedom of signal and noise. The *P*-value of this *F*-ratio is estimated by the cumulative distribution function *F*(*F*_*m*_; *df*_*s*_, *df*_*n*_). A complex *F*-value incorporating the SNR and phase, *θ*_*m*_(*ω*_*s*_), of the signal was obtained by:

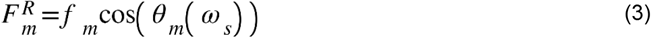

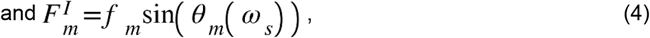

where *f*_*m*_ is the activation amplitude (square root of *F*_*m*_). Each functional dataset was registered with the structural images of subject *S*, and a complex *F*-value was displayed on vertex *v* on the cortical surface of subject *S*, yielding a phase-encoded activation map 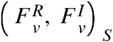 for a single functional scan (Fig. 1A).

The complex *F*-values in voxel *m* were further averaged (voxel-wise) across two scans, *k* ={1, 2}, for the same task within subject *S* by:

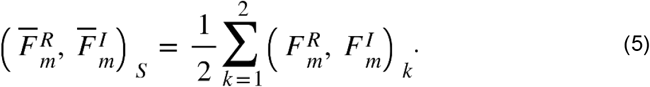

The average dataset was computed using the “Combine 3D Phase Statistics” function in *csurf*. The resulting single-subject vector-average values were then projected onto vertex *v* on the cortical surfaces of subject *S*, yielding a map of 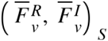.

Surface-based group-average maps for each task were obtained using the spherical averaging method (“Cross Session Spherical Average” function in *csurf*) comprising three steps. First, the vector-average map of subject *S*, 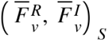, was resampled to a common spherical coordinate system using the *FreeSurfer mri_surf2surf* command (https://freesurfer.net/fswiki/mri_surf2surf), resulting in a new map 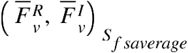 on the cortical surface of *fsaverage*. Second, the complex *F*-values of each vertex *v* on the common spherical surface were vector-averaged (vertex-wise) across *N* subjects by:

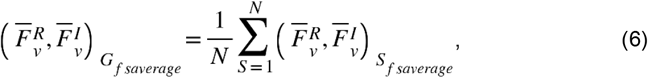

which yielded a group-average map for each task. The amplitude and phase of each vertex in this map were obtained by:

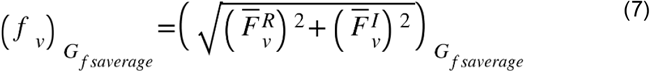

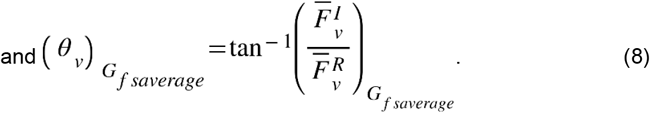

Vertices with significant activations (*F*(2,230) > 4.7, *P* < 0.01) in the map of 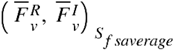 were further tested across subjects (*n* =31, *F*(2,60) > 4.97, *P* < 0.01), and corrected for multiple comparisons using surface-based cluster-size exclusion (cluster = 64 mm^2^, *P* < 0.05, corrected). Finally, the phase and amplitude of each vertex in the group-average maps were displayed on the standard *fsaverage* surface included in *FreeSurfer* (Fig. 1E to J).

#### Analysis in surface-based regions of interest (sROIs)

To illustrate periodic activations (16 cycles per scan) with different phases, we outlined five sROIs from the left hemisphere of Subject 1 (Fig. 1A). In the 4D dataset of an English reading-reciting task for Subject 1, voxels enclosed in each sROI label were identified using the *csurf* command *write_label_timecourses_stats*. The time courses were extracted from all voxels within each sROI and then averaged (Fig. 1B). The complex *F*-value of *V* vertices in each sROI was vector-averaged (vertex-wise) by:

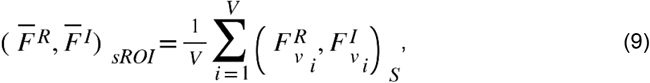

which was displayed as a phasor in the phasor diagram (Fig. 1C). The mean phase of each sROI was obtained by:

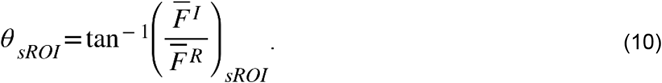

To analyze the activation amplitudes and phases in selected sROIs in the single-subject maps 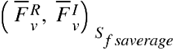 and the group-average map 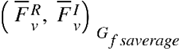, all displayed on the surface of *fsaverage*, we selected task-positive regions for each task (excluding DMN regions Fig. 1E-J), and outlined sROIs including four DMN core regions (vmPFC, AG, PCC, and dmPFC) and one orofacial region (MI/SI) (Fig. 1E-J, cyan contours). For a Task region or an sROI containing *V* vertices in the map of Subject *S* displayed on the *fsaverage* surface, an average complex *F*-value (phasor) was obtained by:

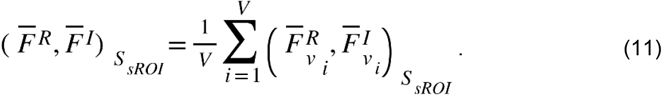

The average amplitude and phase in the Task region or sROI were obtained by:

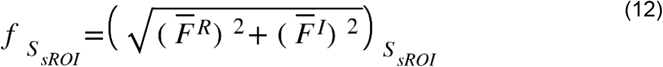

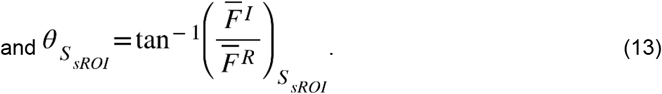

In the phasor diagram, the average complex *F*-value of each Task region or sROI in the single-subject maps was displayed as a colored dot (Fig. 2A). For each task, the distribution of amplitudes (*f*_*S_sROI*_) across subjects is shown in Fig. 2B, and the phase of each sROI is summarized for each subject in Datatset S1.

For a Task region or an sROI containing *V* vertices in the group-average map displayed on the *fsaverage* surface, an average complex *F*-value was obtained by:

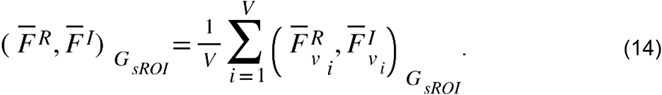

The average amplitude and phase in the Task region or sROI were obtained by:

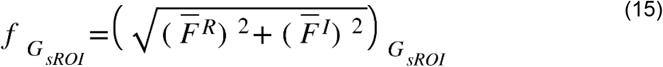

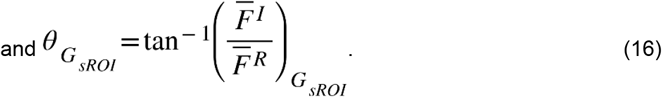

In the phasor diagram, the average complex *F*-value in each Task region or sROI in the group-average maps is displayed as a colored lollipop (Fig. 2A). For each task, the average phase in each Task region or sROI is summarized in Datatset S1.

#### Dataset S1

Average phase (in degree) in each Task region or sROI in the group-average and single-subject maps.

Dataset S1 is available on the Open Science Framework (https://osf.io/hea2p/?view_only=34d7f3c5a37341e9813e04796b0a25d3).

